# Tryptophan catabolism drives AhR-dependent macropinocytosis and tumorigenesis in tuberous sclerosis complex

**DOI:** 10.64898/2026.11.12.765182

**Authors:** Maria Moyet, Aidan McGrath-Conwell, Eleanore Gray, Suzanne Jalbert Jones, Sarah Lafleur, Carleigh Rosenberg, Vishva Patel, James Gray, Windrie Cox, Elizabeth P. Henske, Harilaos Filippakis

**Author notes:** Corresponding author: Harilaos Filippakis, Ph.D. Room 207, Pickus Research Building, University of New England, 11 Hills Beach Road, Biddeford, ME 0400, **Conflict of interest statement:** The authors declare no potential conflicts of interest. Authors contributed equally (Maria Moyet, Aidan McGrath-Conwell).

## Abstract

Tuberous sclerosis complex (TSC) is a rare genetic disorder characterized by hamartomatous tumors of the brain, skin, kidney and lung (lymphangioleiomyomatosis, LAM). TSC and LAM arise from biallelic inactivation of *TSC1* or *TSC2* which drives metabolic reprogramming, including macropinocytic scavenging of extracellular nutrients. To determine which nutrients sustain macropinocytosis, we measured dextran uptake after repletion of individual essential amino acids. Unexpectedly, only tryptophan (Trp) stimulated uptake in TSC2-deficient cells by 2.8-fold compared to amino acid-free medium (*p* < 0.0001), with no effect in TSC2-expressing cells. Steady-state metabolomics revealed that Trp supplementation raised intracellular Trp by 3.5-fold and lowered kynurenine by 60%. IDO1 inhibition abolished the response, indicating that Trp catabolism rather than Trp itself is the proximal signal. Interestingly, *Ido1*, *Tdo2* and *AhR* were markedly upregulated in TSC2-deficient cells, and rapamycin repressed *Ido1* and *Tdo2* but not *AhR*. AhR was constitutively nuclear in TSC2-deficient cells, and rapamycin or the AhR antagonist CH-223191 returned it to the cytoplasm. Inhibition of IDO1, (Linrodostat), TDO2 (LM10) or AhR (StemRegenin-1; SR1, CH-223191) selectively decreased macropinocytosis and proliferation of TSC2-deficient cells. AhR antagonism also depleted glutathione by 70%. Treatment of mice with bearing TSC2-deficient tumors with SR1 or CH-223191 reduced tumor volume by approximately 70% and 90% (*p* < 0.0001), respectively. Both treatments also strongly decreased mTORC1 activity in tumors, consistent with a model in which macropinocytosis drives mTOR activity in TSC. Collectively, our data identify a Trp-kynurenine-AhR axis that supports macropinocytosis in TSC2-deficient cells, and suggest AhR antagonists as a novel therapeutic approach for the treatment of TSC and LAM.

**Statement of Significance:** We show that mTORC1 hyperactivation drives tryptophan catabolism to kynurenine, which activates AhR and sustains macropinocytic nutrient scavenging in TSC2-deficient cells. AhR antagonists suppress tumor growth alone, and more thoroughly with rapamycin. This work identifies a novel therapeutic pathway for patients with TSC and LAM.

## Introduction

Tuberous sclerosis complex (TSC) is a rare autosomal dominant genetic disease caused by loss-of-function mutations in either *TSC1* or *TSC2* and affects approximately 1 in 6,000 individuals worldwide, producing a broad spectrum of hamartomatous tumors across multiple organ systems including the brain, kidney, heart, skin, and lung (1,2). The pulmonary manifestation of TSC, lymphangioleiomyomatosis (LAM), is a progressive, destructive cystic lung disease occurring almost exclusively in women and is characterized by the infiltration of neoplastic smooth muscle-like cells that destroy alveolar architecture, leading to respiratory failure in a substantial number of affected individuals (3,4). Both sporadic and TSC-associated LAM arise through biallelic loss of *TSC2*, typically a germline or somatic mutation followed by loss of heterozygosity, which leads to complete loss of the TSC2 tumor suppressor protein tuberin and constitutive hyperactivation of the mechanistic target of rapamycin complex 1 (mTORC1) (1,5,6). The rapalog sirolimus is the only FDA-approved therapy for LAM and TSC-associated tumors, and while it produces meaningful clinical benefit, stabilizing lung function and reducing tumor burden, its effects are cytostatic, and disease progression invariably recurs upon cessation of treatment (7,8). The genomic landscape of TSC reveals the full spectrum of *TSC1* and *TSC2* loss-of-function mutations across missense, frameshift, and splice-site variants (9), yet all converge on constitutive mTORC1 hyperactivation as the central pathogenic mechanism. This clinical limitation underscores the need to identify novel dependencies in TSC2-deficient cells that can be targeted to achieve durable tumor suppression.

mTORC1 is a master regulator of anabolic cell growth that coordinates a metabolic transcriptional program to support the biosynthesis of proteins, nucleotides, and lipids required for cell proliferation(10). In TSC, constitutive mTORC1 hyperactivation creates biosynthetic demands that cannot be fully met through endogenous biosynthetic pathways, creating a dependency on exogenous nutrient sources (11). For example, TSC2-deficient cells are dependent on glucose and glutamine and must utilize multiple nutrient scavenging strategies to maintain metabolic homeostasis, including lysosome-dependent acquisition of extracellular cholesterol (12).

Macropinocytosis is an evolutionarily conserved, actin-dependent form of endocytosis in which plasma membrane ruffles close into macropinocytic cups that engulf extracellular fluid and macromolecules in bulk. The resulting macropinosomes traffic to lysosomes, where internalized proteins are degraded into amino acids and other nutrients (13–15). Landmark studies in Ras-transformed pancreatic ductal adenocarcinoma cells established macropinocytosis as a critical amino acid supply route in the nutrient-poor tumor microenvironment (16), and subsequent work has demonstrated that this process is reciprocally coupled to mTORC1 signaling in cancer cells (17). We have previously demonstrated that macropinocytosis is markedly elevated in TSC2-deficient cells and that pharmacologic or genetic inhibition of macropinocytic uptake selectively impairs their proliferation and tumorigenesis (18). Our prior work identified the lipid kinase diacylglycerol kinase alpha (DGKA) as a key mechanistic driver of macropinocytosis in TSC, and demonstrated that ritanserin, an inhibitor of DGKA, rewires phospholipid metabolism in TSC2-deficient cells and suppresses tumor formation in preclinical models (19). This body of evidence establishes macropinocytosis as a therapeutic vulnerability in TSC; however, the upstream signals that initiate and sustain elevated macropinocytosis in TSC2-deficient cells remain incompletely defined.

Among the essential amino acids, tryptophan (Trp) occupies a unique position at the intersection of nutrient sensing, metabolic reprogramming, and oncogenic signaling. In addition to its incorporation into proteins, Trp serves as the precursor for a diverse set of bioactive metabolites that influence cellular physiology through multiple mechanisms (20,21). The predominant catabolic route for Trp in most cell types is the kynurenine pathway, which is initiated by the rate-limiting enzymes indoleamine 2,3-dioxygenase 1 (IDO1) and tryptophan 2,3-dioxygenase (TDO2). Together, IDO1 and TDO2 convert Trp to N-formylkynurenine, which is rapidly hydrolyzed to kynurenine (Kyn). Kynurenine and several of its downstream metabolites are high-affinity endogenous ligands for the aryl hydrocarbon receptor (AhR), a ligand-activated basic helix-loop-helix/Per-ARNT-Sim (bHLH-PAS) transcription factor. Upon ligand binding, AhR translocates to the nucleus, heterodimerizes with the aryl hydrocarbon receptor nuclear translocator (ARNT), and drives the transcription of a broad set of target genes (22). The IDO1–kynurenine–AhR axis has been most thoroughly characterized in the context of tumor immune evasion, where it generates local Trp depletion that impairs effector T cell function and promotes the differentiation of immunosuppressive regulatory T cells, thereby enabling tumor cells to evade antitumor immunity (23–25). Beyond these immunologic functions, emerging evidence indicates that the kynurenine–AhR pathway also operates in a cell-autonomous manner within tumor cells to support their survival and proliferation (26). IDO1 has been reported to inhibit Trp-dependent stimulation of mTOR, establishing a functional link between Trp catabolism and mTORC1 status (27). However, the role of mTORC1 hyperactivation on Trp catabolism has not been investigated in TSC. We therefore hypothesized that constitutive mTORC1 activity in TSC upregulates IDO1 and TDO2 expression, leading to increased kynurenine production, constitutive AhR nuclear translocation, and enhanced macropinocytic nutrient uptake.

Here, we demonstrate that extracellular Trp is a potent stimulator of macropinocytosis in TSC2-deficient cells and that this stimulation is dependent on kynurenine pathway activity and downstream AhR signaling. We further demonstrate that AhR is localized to the nucleus of TSC2-deficient cells, and that pharmacologic blockage of IDO1, TDO2, or AhR selectively impairs macropinocytosis and proliferation of TSC2-deficient cells. Additionally, our data show that AhR inhibition depletes glutamate and glutathione stores in TSC2-deficient cells. Finally, we provide preclinical evidence that targeting the kynurenine–AhR axis with pharmacologic agents, alone or in combination with rapamycin, suppresses TSC2-deficient tumor growth *in vivo*. These findings define the Trp–Kyn–AhR–macropinocytosis axis as a cell-autonomous and targetable metabolic vulnerability in TSC, with direct therapeutic implications for patients with TSC and LAM.

## Materials and Methods

### Cell lines and cell culture

Littermate-derived *Tsc2*^−/−^*Tp53*^−/−^ and *Tsc2*^+/+^*Tp53*^−/−^ mouse embryonic fibroblasts (MEF) were provided by David J. Kwiatkowski (Brigham and Women’s Hospital, Boston, MA. 105K is a cell line derived from a *Tsc2*^+/−^ C57BL/6 mouse renal tumor and confirmed to have lost the second *Tsc2* allele, to lack Tsc2 expression and to have hyperactive mTORC1. 105K+TSC2 and 105K-EV cells were generated by re-expressing TSC2 in 105K cells by retroviral delivery of the pLXIN-IRES-hygromycin vector carrying full-length human *TSC2* or empty vector, respectively (12). Unless stated otherwise, cells were cultured at 37°C in 5% CO₂ in Dulbecco’s modified Eagle medium (DMEM) containing 4.5 g/L glucose supplemented with 10% FBS, 100 U/mL penicillin and 100 µg/mL streptomycin. All cells tested negative for *Mycoplasma* and were re-tested bi-monthly using MycoAlert (Lonza). Experiments were performed on cells within 10 passages of thawing.

### Antibodies, drugs and reagents

Rapamycin was purchased from LC Laboratories. 5-(*N*-ethyl-*N*-isopropyl)amiloride (EIPA), L-tryptophan (T0254) and L-kynurenine (K8625) were purchased from Sigma-Aldrich. Linrodostat (BMS-986205) (MedChemExpress, HY-101560), LM10 (MedChemExpress, HY-33298), StemRegenin 1 (MedChemExpress, HY-15001), CH-223191 (MedChemExpress, HY-12684) and JPH-203 were purchased from. AhR was detected with Proteintech antibody 28727-1-AP (1:1000). Compounds, nominal targets, working concentrations and vehicles are listed in Supplementary Table S1. Stock solutions were prepared in dimethyl sulfoxide (DMSO) unless stated otherwise, stored at −20°C protected from light and diluted into culture medium immediately before use; the final DMSO concentration did not exceed 0.1% (v/v) and vehicle- only controls were included in every experiment.

### Amino acid deprivation and repletion

Cells were seeded in 6-well plates and grown in DMEM containing 1% FBS overnight. Cells were washed twice with PBS and incubated overnight in amino acid-free DMEM (US Biological, D9800-27) supplemented with 1% dialyzed FBS (Gibco, 26400044). Amino acid-free medium was then replaced with amino acid-free medium repleted with either the complete set of nine essential amino acids (Gibco, 11130051) or a single essential amino acid, each at the concentration found in DMEM. Trp was used at 68 µM and kynurenine at 100 µM. Repletion was for 24 hours before addition of FITC-dextran or harvest for metabolite profiling.

### Macropinocytosis uptake assays and flow cytometry

Macropinocytosis was assessed as described previously (28,29). Cells were seeded in 6-well plates and grown in 1% FBS DMEM overnight. Treatment with EIPA (25 µM), linrodostat 20 µM, SR-1 (2 µM), CH-223191 (10 µM) or JPH-203 (100 or 200 µM) was carried out for 16 hours, followed by addition of 0.5 mg/mL 70 kDa FITC-dextran (Invitrogen) for 1 hour. Cells were washed twice with ice-cold PBS, trypsinized, recovered in 2% serum phenol red-free DMEM and centrifuged at 425 × *g* for 2 minutes. The pelleted cells were resuspended in 200μL of 2% parafolmadehyde and incubated for 10 minutes, and centrifuged at 425g × *g* for 2 minutes. The supernatant was removed and cells were then resuspended 300 µL of serum-free, phenol red-free DMEM and kept on ice. Fluorescence was assessed by flow cytometry (BD Accuri C6) and analyzed with FlowJo (TreeStar). Median fluorescence intensity of FITC was measured and reported for each sample.

### Microscopy

Cells were seeded on 4-chamber tissue culture glass slides in 1% FBS DMEM overnight, treated with inhibitors and incubated with FITC-dextran as described above. Cells were rinsed twice with PBS and fixed with 4% paraformaldehyde. Confocal images were captured on a FluoView FV-10i Olympus laser point scanning confocal microscope using a 60× objective. Confocal filters (excitation/emission, nm) were 358/461 (DAPI) and 494/521 (FITC-dextran). Epifluorescence images were captured on a Keyence BZ-X710 fluorescent microscope using 40x objective. Acquisition settings were held constant across conditions within each experiment.

### Crystal violet proliferation assay

Cells were seeded at 1,500 cells per well in 96-well plates and allowed to attach overnight. Treatments were administered the following day. After 24, 48 or 72 hours, cells were fixed with 10% formalin for 10 minutes, stained with 0.05% (w/v) crystal violet in distilled water for 30 minutes, washed three times by submerging the plates in clean tap water and air dried. Crystal violet was solubilized by adding 100 µL of methanol per well and absorbance was measured on a plate reader (OD540; Molecular Devices SpectraMax i3).

### mRNA expression analysis

RNA extraction was performed using the RNeasy Micro Kit (Qiagen). Two micrograms of total RNA were reverse transcribed using the High-Capacity cDNA Reverse Transcription Kit (Applied Biosystems). Forty nanograms of cDNA were used for qPCR reactions with TaqMan probes and TaqMan Fast Advanced Master Mix (Applied Biosystems) on an Applied Biosystems instrument. Data were analyzed using the ΔΔCt method, with actin as the reference gene. TaqMan assay identifiers were: *AhR*; Mm00478932_m1, *Ido1;* Mm00492590_m1, *Tdo2;* Mm00451269_m1*, Actb;* Mm02619580_g1.

### Immunofluorescence and analysis of AhR subcellular localization

Cells were seeded on 4-chamber tissue culture glass slides in 1% FBS DMEM overnight and treated with rapamycin (20 nM) or CH-223191 (10 µM) for 24 hours. Cells were rinsed twice with PBS, fixed in 4% paraformaldehyde for 10 minutes, permeabilized in 0.2% Triton X-100 in 1X PBS for 15 minutes, blocked in 2% BSA in 1X PBS and incubated with anti-AhR antibody (1:250), followed by AlexaFluor 488 (Thermofisher; A-11094). Nuclei were counterstained with DAPI and slides were mounted in. Images were captured on a FluoView FV-10i Olympus laser point scanning confocal microscope using a 60× objective, with acquisition settings held constant across conditions.

### Steady-state metabolite profiling by LC-MS/MS

Metabolites from four replicate 60-mm plates per condition were extracted with 80% aqueous methanol. Metabolites were processed using selected reaction monitoring (SRM) with polarity switching on a 5500 QTRAP triple quadrupole mass spectrometer (AB/SCIEX) coupled to a Prominence UFLC HPLC system (Shimadzu) using amide HILIC chromatography (Waters) at pH 9.2 (BIDMC Mass Spectrometry Core, Beth Israel Deaconess Medical Center, Boston, MA). Two hundred fifty-two endogenous water-soluble metabolites were measured at steady state. Peak area data were normalized to the protein concentration of three additional replicate plates per condition. Cells were harvested 24 hours after Trp repletion and SR-1 treatment.

### Metabolite set enrichment analysis

Normalized peak area data were uploaded to MetaboAnalyst v. 6.0 (http://www.metaboanalyst.ca) for processing and metabolite set enrichment analysis. Data were filtered by interquartile range and auto scaled (mean-centered and divided by the standard deviation of each variable), and log₂ transformation was applied. For two-group comparisons, a *t* test followed by false discovery rate correction was used; two-way ANOVA was applied for comparisons of more than two groups. Significance was defined as *q* < 0.05 and *p* < 0.05, respectively.

### Animal studies

All animal studies were performed in accordance with institutional protocols approved by the University of New England Institutional Animal Care and Use Committee. Xenografts were generated by subcutaneously injecting 2.5 × 10⁶ 105K cells mixed 1:1 (v/v) with Matrigel (Corning, 356237) to a final volume of 100–150 µL unilaterally into the shoulders of anesthetized mice using a 21 G needle. 8-week-old male and female C57BL/J6 mice were used. Tumors were measured every two days by caliper once palpable, and tumor volume was calculated as (length × width²)/2. When mean tumor volume reached 100 mm³, mice were randomized to treatment arms of *n* = 8 per group. In the first study, mice received vehicle, SR-1 (6mg/kg intraperitoneally, Monday to Friday) or rapamycin (3 mg/kg intraperitoneally, Monday, Wednesday and Friday). In the second study, mice received vehicle, CH-223191 (10 mg/kg, Monday to Friday), Linrodostat (10 mg/kg, Monday to Friday), rapamycin (3 mg/kg, Monday, Wednesday and Friday), or rapamycin combined with CH-223191 at the same doses and schedules. Treatments were administered intraperitoneally and 0.25% PEG200, 0.25% Tween-80 in water was used as vehicle. Treatment continued for 38 days and mice were euthanized 24 hours after the final dose. No animals were excluded from analysis.

### Immunohistochemistry

Excised tumors were fixed in 10% neutral buffered formalin for 24 hours, embedded in paraffin and sectioned at 5 µm. Sections were deparaffinized, rehydrated and subjected to heat-induced epitope retrieval. Endogenous peroxidase activity was quenched and sections were blocked. Sections were incubated with antibodies against Ki-67 Cell Signaling; 12202), phospho-p70 S6 kinase (Cell Signaling # 8753, 1:250). Sections were counterstained with hematoxylin and imaged on an Olympus VS200 using an 20x objective.

### Statistical analysis

Normally distributed data were analyzed for statistical significance with Student’s unpaired *t* test, and multiple comparisons were made with one-way and two-way ANOVA with Bonferroni correction. Tumor growth curves (Fig. 6A and C) were analyzed by two-way ANOVA with treatment and time as factors, followed by Dunnett’s multiple comparisons test against the vehicle group at the final time point. Two pre-specified comparisons in Fig. 6C — rapamycin plus CH-223191 versus rapamycin alone, and versus CH-223191 alone — were made at the final time point from the same ANOVA, with Bonferroni correction for the two tests.

### Data availability

The metabolomics data generated in this study are available on request from the corresponding author. All other data generated in this study are available within the article and its supplementary data files.

## Results

### Tryptophan selectively stimulates macropinocytosis in TSC2-deficient cells

To investigate whether specific amino acids differentially regulate macropinocytosis in TSC2-deficient versus TSC2-expressing cells, we assessed uptake of 70 kDa FITC-dextran, a well-validated macropinocytosis marker, in *Tsc2*^−/−^ and *Tsc2*^+/+^ mouse embryonic fibroblasts (MEFs) under defined amino acid conditions. Cells were maintained in DMEM containing 1% FBS and subjected to amino acid-free (aaf) conditions, followed by repletion with either the complete set of essential amino acids (eaa) or individual essential amino acids. Fluorescence microscopy revealed an accumulation of punctate cytoplasmic dextran fluorescence in *Tsc2*^−/−^ MEFs supplemented with tryptophan (Trp; 68 µM), reflecting the characteristic vesicular pattern of macropinocytic uptake (Fig. 1A). In contrast, *Tsc2*^+/+^ MEFs supplemented with Trp under identical conditions showed no increase in macropinocytosis compared with amino acid-free controls, indicating that the macropinocytic response to Trp requires TSC2 loss (Fig. 1A). Treatment with the macropinocytosis inhibitor 5-(N-ethyl-N-isopropyl)amiloride (EIPA; 25 µM) abrogated Trp-stimulated dextran uptake in *Tsc2*^−/−^ cells, consistent with macropinocytic internalization rather than receptor-mediated endocytosis or fluid-phase pinocytosis (Fig. 1A). To quantify the macropinocytic response systematically, we performed flow cytometric analysis of dextran uptake across amino acid conditions in *Tsc2*^−/−^ MEFs. Supplementation with a complete mixture of essential amino acids increased dextran uptake by approximately 55% compared with amino acid-free baseline (p < 0.01), and this effect was reversed by EIPA, which reduced uptake to approximately 30% below the amino acid-free baseline (p< 0.0001; Fig. 1B). To identify the responsible amino acid(s), we tested each essential amino acid individually. Of the essential amino acids tested, Trp was the only one that induced macropinocytosis: Trp supplementation alone increased dextran uptake approximately 2.8-fold relative to amino acid-free medium (p< 0.0001; Fig. 1C). Trp add-back alone restored uptake to approximately 85% of the level measured in complete 1% FBS medium, whereas none of the other eight essential amino acids differed from amino acid-free medium (Fig. 1C). EIPA reduced Trp-stimulated uptake by approximately 50%, to a level that remained marginally above the amino acid-free baseline (p< 0.05 versus amino acid-free), showing that macropinocytosis is the primary uptake mechanism (Fig. 1C). These results demonstrate that Trp exerts a selective effect on macropinocytic activity in TSC2-deficient cells that is not shared by other essential amino acids.

**Figure 1:**
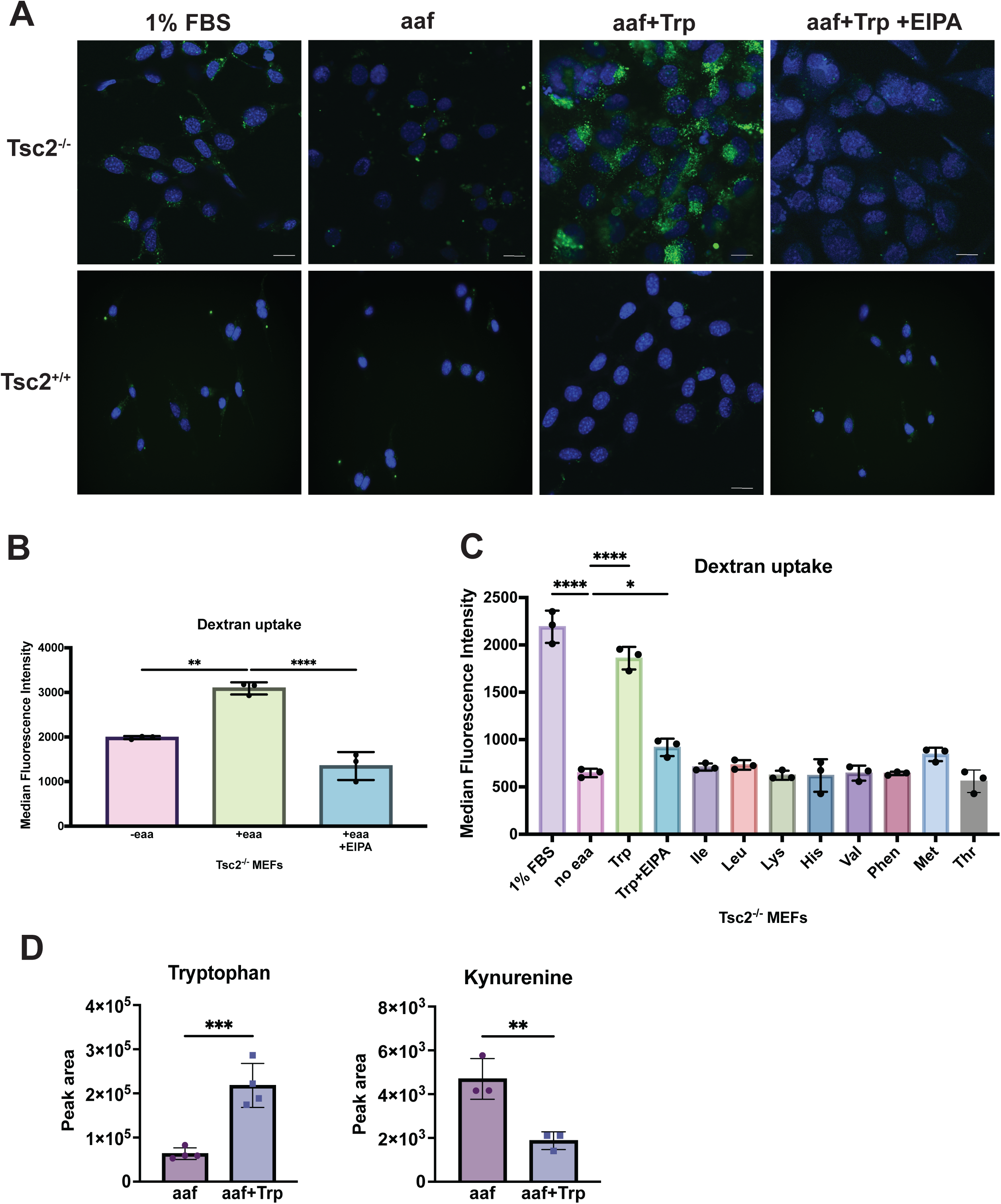
Tryptophan enhances macropinocytosis in TSC2-deficient cells. **(A)** TSC2-deficient (*Tsc2*⁻/⁻) and TSC2-expressing (*Tsc2*⁺/⁺) mouse embryonic fibroblasts (MEFs) were grown in DMEM containing 1% FBS, in amino acid-free medium (aaf), or in aaf medium repleted with tryptophan (Trp; 68 µM) for 24 h, with or without 5-(N-ethyl-N-isopropyl)amiloride (EIPA; 25 µM, 16 h), and then incubated with 0.5 mg/mL 70 kDa FITC-dextran (green) for 1 h. Nuclei were counterstained with DAPI (blue). Trp increased punctate cytoplasmic dextran uptake in *Tsc2*⁻/⁻ but not *Tsc2*⁺/⁺ MEFs, and EIPA abolished the response. Representative confocal images acquired with identical settings across conditions. Scale bar, 100µm. **(B)** Flow cytometric quantification of dextran uptake in *Tsc2*⁻/⁻ MEFs. Repletion with the complete set of nine essential amino acids (eaa) increased median fluorescence intensity by approximately 55% relative to the amino acid-free baseline (p < 0.01), and EIPA reduced uptake to approximately 30% below that baseline (p < 0.0001). **(C)** Repletion with individual essential amino acids. Trp alone increased dextran uptake approximately 2.8-fold relative to amino acid-free medium (p < 0.0001), restoring uptake to approximately 85% of the level measured in complete 1% FBS medium; EIPA reduced Trp-stimulated uptake by approximately 50%, to a level that remained marginally above the amino acid-free baseline (p < 0.05 versus amino acid-free). None of the other eight essential amino acids differed from amino acid-free medium. **(D)** Steady-state metabolite profiling by LC-MS/MS of *Tsc2*⁻/⁻ MEFs harvested 24 h after Trp repletion. Intracellular Trp increased approximately 3.5-fold (p < 0.001) and kynurenine decreased by approximately 60% (p < 0.01). A–C, n = 3 biological replicates; D, n = 4 biological replicates. Data are mean ± SD. B and C were analyzed by two-way ANOVA with Bonferroni correction and D by unpaired two-tailed Student t test. *, p < 0.05;, p < 0.01; *, P < 0.001, P < 0.0001.

To determine whether the macropinocytic effect of Trp requires its enzymatic conversion to kynurenine intracellularly, we asked whether direct supplementation with Kyn could substitute for Trp. Kyn supplementation failed to increase dextran uptake above amino acid-free baseline levels in *Tsc2*^−/−^ MEFs (p = ns; Supplementary Fig. S1A), and although co-treatment with linrodostat produced no further change, EIPA reduced uptake in Kyn-supplemented cells by approximately 70% (p< 0.0001), to the same level as in amino acid-free cells treated with EIPA (Supplementary Fig. S1A). Extracellular kynurenine is therefore not sufficient to drive macropinocytosis. In parallel, Linrodostat abolished the Trp-stimulated increase in dextran uptake, returning it to a level indistinguishable from amino acid-free medium (p= ns versus amino acid-free), whereas EIPA reduced it by approximately 80% (p< 0.0001; Supplementary Fig. S1A), indicating that IDO1-dependent Trp catabolism is required for the macropinocytic response to Trp. Fluorescence imaging confirmed the absence of increased dextran-positive macropinosomes following Kyn supplementation in both cell lines, in contrast to the vesicular uptake elicited by Trp (Supplementary Fig. S1B). To determine whether the plasma membrane large neutral amino acid transporter LAT1/SLC7A5 mediates the macropinocytic response to Trp, we treated both *Tsc2*^−/−^ and *Tsc2*^+/+^ MEFs with the LAT1 inhibitor JPH-203 (100 µM, 200 µM) or EIPA. Basal dextran uptake was significantly higher in *Tsc2*^−/−^ cells than in *Tsc2*^+/+^ cells (p< 0.0001), confirming our previous findings (9, 10). EIPA significantly reduced uptake in both cell lines (p < 0.01), whereas JPH-203 had no effect on macropinocytosis (Supplementary Fig. S1C). Together, these data implicating intracellular Trp catabolism rather than Trp or Kyn transport as the proximal signal.

Because Trp is the entry substrate for the kynurenine pathway and its catabolism produces AhR-activating metabolites, we characterized the intracellular metabolic response to exogenous Trp in *Tsc2*^−/−^ MEFs. Targeted steady-state metabolite profiling by LC-MS demonstrated that Trp supplementation produced an approximately 3.5-fold increase in intracellular Trp levels (*p* < 0.001; Fig. 1D), confirming cellular uptake of the exogenous amino acid. Intracellular kynurenine levels fell by approximately 60% under the same conditions (*p*< 0.01; Fig. 1D).

### Kynurenine pathway enzymes and Ahr are upregulated in TSC2-deficient cells

To determine the status of the kynurenine pathway in mTORC1 hyperactive cells, we quantified the gene expression of the two rate-limiting enzymes *IDO1,* and *TDO2* together with *AhR*. Quantitative RT-PCR analysis of *Tsc2*^−/−^ and *Tsc2*^+/+^ MEFs revealed that *AhR* transcript levels were elevated approximately 50-fold in TSC2-deficient cells compared with TSC2-expressing controls (*p* < 0.0001; Fig. 2A). *Tdo2* was similarly elevated approximately 45-fold in *Tsc2*^−/−^ MEFs (*p* < 0.0001; Fig. 2B). Rapamycin treatment (20 nM, 24 hours) decreased *Tdo2* expression by approximately 80% in *Tsc2*^−/−^ cells (*p* < 0.0001; Fig. 2B), indicating that mTORC1 activity is required to maintain this elevated expression. Consistent with these findings, *Ido1* transcript levels were increased 7.5-fold in *Tsc2*^−/−^ MEFs relative to *Tsc2*^+/+^ MEFs (*p* < 0.01; Fig. 2C), and rapamycin reduced *Ido1* expression by approximately 80% in TSC2-deficient cells (*p* < 0.01; Fig. 2C). Rapamycin did not produce a comparable change in *Tdo2* or *Ido1* expression in *Tsc2*^+/+^ MEFs, confirming that the mTORC1 sensitivity of these transcriptional changes is specific to the TSC2-deficient context. In contrast to *Ido1* and *Tdo2*, *AhR* transcript levels were not reduced by rapamycin in *Tsc2*^−/−^ MEFs (Fig. 2A). These data establish that the kynurenine pathway genes *Ido1*, *Tdo2* and *AhR* are all upregulated in TSC2-deficient cells and may play a role in facilitating macropinocytosis.

**Figure 2:**
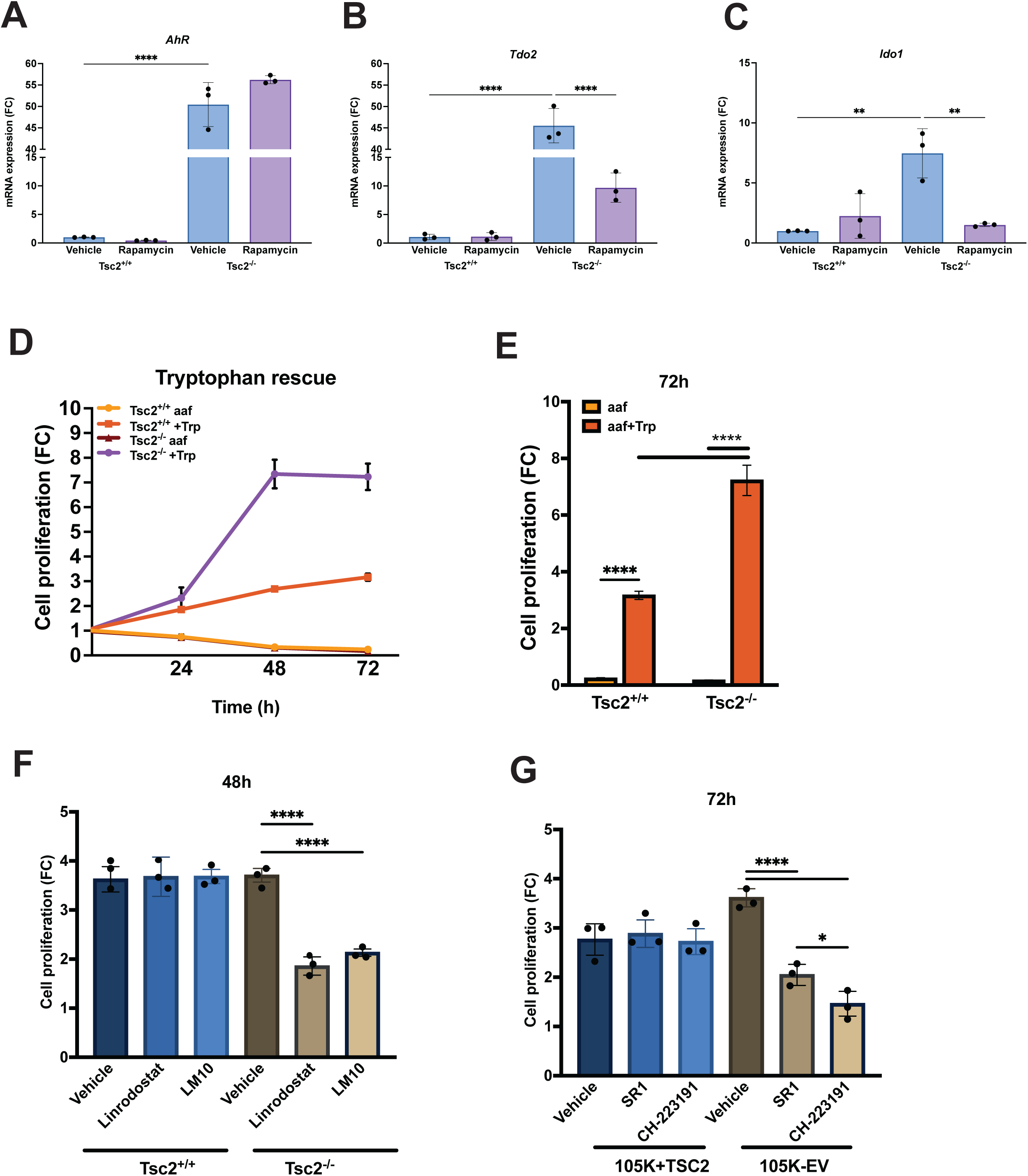
TSC2-deficient cells depend on Trp and the kynurenine pathway for survival. **(A–C)** Quantitative RT-PCR of *Ahr*, *Tdo2* and *Ido1* in *Tsc2*⁻/⁻ and *Tsc2*⁺/⁺ MEFs treated with vehicle or rapamycin (20 nM, 24 h). **(A)** *Ahr* transcript was elevated approximately 50-fold in *Tsc2*⁻/⁻ MEFs (P < 0.0001) and was not reduced by rapamycin. **(B)** *Tdo2* was elevated approximately 45-fold (p < 0.0001) and rapamycin reduced it by approximately 80% in *Tsc2*⁻/⁻ MEFs (p < 0.0001). **(C)** *Ido1* was elevated 7.5-fold (p < 0.01) and rapamycin reduced it by approximately 80% (p < 0.01). Rapamycin did not produce a comparable change in Tdo2 or Ido1 expression in Tsc2 ⁺/⁺ MEFs. **(D)** Crystal violet proliferation of *Tsc2*⁻/⁻ and *Tsc2*⁺/⁺ MEFs in amino acid-free medium with or without Trp (68 µM) over 72 h, expressed as fold change from day 0. Trp increased *Tsc2*⁻/⁻ proliferation 7.2-fold at 72 h (p < 0.0001) compared with 3.1-fold in *Tsc2*⁺/⁺ MEFs. **(E)** The same comparison at 72 h. **(F)** Inhibition of IDO1 with linrodostat (20 µM) or of TDO2 with LM10 (20 µM) reduced *Tsc2*⁻/⁻ proliferation at 48 h by approximately 50% and 40%, respectively (p < 0.0001 for both), with no significant effect on *Tsc2*⁺/⁺ MEFs. **(G)** Inhibition of AhR with SR-1 (2 µM) or CH-223191 (10 µM) reduced proliferation of TSC2-deficient 105K-EV cells at 72 h by approximately 45% and 60%, respectively (p < 0.0001), whereas proliferation of the isogenic TSC2 re-expressing line (105K+TSC2) was not significantly affected. n = 3 biological replicates. Proliferation was quantified by crystal violet staining. Data are mean ± SD; two-way ANOVA with Bonferroni correction. *, p < 0.05; p< 0.01;, p < 0.0001.

We next asked whether this elevated kynurenine pathway activity has functional consequences for TSC2-deficient cell proliferation. Trp supplementation (68 µM, 72 hours) increased *Tsc2*^−/−^ proliferation 7.2-fold compared to day 0 (*p* < 0.0001; Fig. 2D and E), whereas proliferation of *Tsc2*+/+ cells increased only 3.1-fold. To test whether Trp-derived kynurenine pathway metabolites support proliferation of TSC2-deficient cells, we treated *Tsc2*^−/−^ and *Tsc2*^+/+^ MEFs with the selective IDO1 inhibitor linrodostat (BMS-986205; 20 µM) or the selective TDO2 inhibitor LM10 (20 µM). Linrodostat and LM10 reduced *Tsc2*^−/−^ proliferation by approximately 50% and 40%, respectively, at 48 hours (*p* < 0.0001 for both), whereas neither compound significantly affected proliferation of *Tsc2*^+/+^ MEFs (Fig. 2F), indicating that both IDO1 and TDO2 contribute to the proliferative capacity of TSC2-deficient cells. We next examined the downstream AhR node directly. Treatment of the TSC2-deficient mouse kidney tumor-derived cell line 105K (105K-EV, empty vector) and its isogenic TSC2 re-expressing derivative (105K+TSC2) with the AhR antagonists SR-1 (2 µM) or CH-223191 (10 µM) decreased 105K-EV proliferation by approximately 45% and 60%, respectively (*p* < 0.0001; Fig. 2G), whereas proliferation of 105K+TSC2 cells was not significantly affected by either inhibitor at these concentrations (Fig. 2G). These data indicate that mTORC1-driven upregulation of the kynurenine pathway and downstream AhR activation are required for the proliferation of TSC2-deficient cells.

### AhR inhibition suppresses macropinocytosis in TSC2-deficient cells

We next sought to determine whether IDO1–kynurenine–AhR signaling is mechanistically required for macropinocytosis in TSC. Fluorescence imaging of *Tsc2*^−/−^ MEFs treated with Linrodostat, SR-1, or EIPA showed reduced dextran fluorescence puncta compared with vehicle-treated *Tsc2*^−/−^ controls (Fig. 3A). To quantify the extent of macropinocytosis inhibition we performed flow cytometry. Interestingly, Linrodostat treatment reduced macropinocytic uptake by approximately 30% (*p* < 0.0001; Fig. 3B), whereas SR-1 produced a 70% reduction and EIPA an 89% reduction (*p* < 0.0001; Fig. 3B). These results indicate that IDO1 inhibition alone provides only partial suppression of macropinocytosis and point to AhR as the principal node through which the pathway controls macropinocytic uptake in TSC.

**Figure 3:**
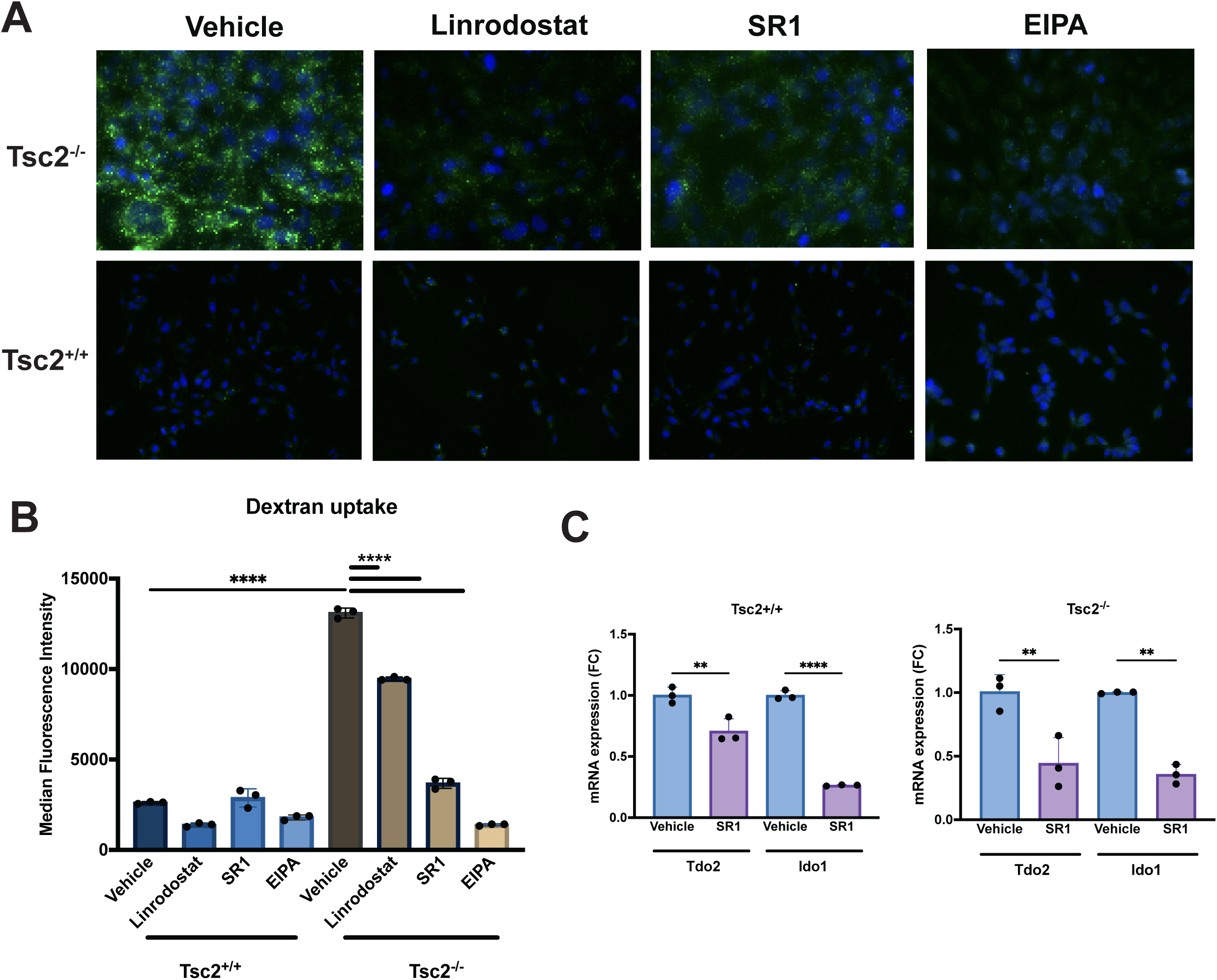
Kynurenine pathway inhibition blocks macropinocytosis in TSC2-deficient cells. **(A)** *Tsc2*⁻/⁻ and *Tsc2*⁺/⁺ MEFs were treated with vehicle, linrodostat (20 µM), SR-1 (2 µM) or EIPA (25 µM) for 16 h and then incubated with 0.5 mg/mL 70 kDa FITC-dextran (green) for 1 h; nuclei were counterstained with DAPI (blue). Each inhibitor reduced dextran-positive puncta in *Tsc2*⁻/⁻ cells. Representative epifluorescence. images acquired with identical settings. Scale bar, 100µm. **(B)** Flow cytometric quantification. Basal dextran uptake was higher in *Tsc2*⁻/⁻ than in *Tsc2*⁺/⁺ MEFs (p < 0.0001). Relative to *Tsc2*⁻/⁻ vehicle, linrodostat reduced uptake by approximately 30%, SR-1 by approximately 70% and EIPA by approximately 89% (p < 0.0001 for all). **(C)** SR-1 (2 µM, 24 h) decreased *Tdo2* and *Ido1* transcript levels in *Tsc2*⁻/⁻ MEFs (−57% and −66%) and in *Tsc2*⁺/⁺ MEFs (−29% and −74%). n = 3 biological replicates. Data are mean ± SD; two-way ANOVA with Bonferroni correction. **\*\*,** p < 0.01; ****, p < 0.0001.

We also found that SR-1 treatment decreased the expression of both *Tdo2* and *Ido1* in *Tsc2*^−/−^ MEFs (*Tdo2*, −57%; *Ido1*, −66%) and, to a similar extent, in *Tsc2*^+/+^ MEFs (*Tdo2*, −29%; *Ido1*, −74%) (Fig. 3C), consistent with an AhR-dependent positive feedback loop in which AhR activity promotes expression of the enzymes that generate its ligand. This type of feedback has been described for AhR in immune cell contexts (21, 22) but has not previously been characterized in TSC2-deficient cells. These observations suggest that AhR antagonism acts at two levels, blocking downstream signaling and reducing expression of the upstream enzymes that sustain ligand production, which may account for its greater effect on macropinocytosis relative to IDO1 inhibition.

### AhR is localized to the nucleus of TSC2-deficient cells in an mTORC1-dependent manner

Canonical AhR signaling is initiated by ligand binding in the cytoplasm, which promotes conformational changes enabling nuclear import, heterodimerization with ARNT, and transcriptional activation of target genes containing xenobiotic/dioxin response elements (XRE/DRE). To determine whether kynurenine pathway activity in TSC2-deficient cells is sufficient to drive constitutive AhR nuclear translocation, we examined AhR subcellular localization in *Tsc2*^−/−^ and *Tsc2*^+/+^ MEFs. AhR co-localized extensively with nuclear DAPI staining in *Tsc2*^−/−^ MEFs, indicating constitutive nuclear AhR localization (Fig. 4). In contrast, AhR in *Tsc2*^+/+^ MEFs displayed a predominantly cytoplasmic distribution, consistent with the quiescent state of the receptor in the absence of exogenous ligand or mTORC1 hyperactivation (Fig. 4). Rapamycin redistributed AhR from nuclear to cytoplasmic compartments in *Tsc2*^−/−^ cells, indicating that constitutive mTORC1 activity is required to sustain the nuclear AhR pool, potentially through its effect on *Ido1* and *Tdo2* expression (Fig. 4). Consistent with a ligand-driven mechanism, treatment with CH-223191 (10 µM, 24 hours) similarly blocked nuclear AhR localization in *Tsc2*^−/−^ MEFs (Fig. 4). Neither rapamycin nor CH-223191 altered AhR localization in *Tsc2*^+/+^ MEFs, in which AhR remained cytoplasmic under all conditions examined. These data indicate that nuclear AhR in TSC2-deficient cells is mTORC1-dependent, and reflects cell-autonomous activation of this pathway following TSC2 loss.

**Figure 4:**
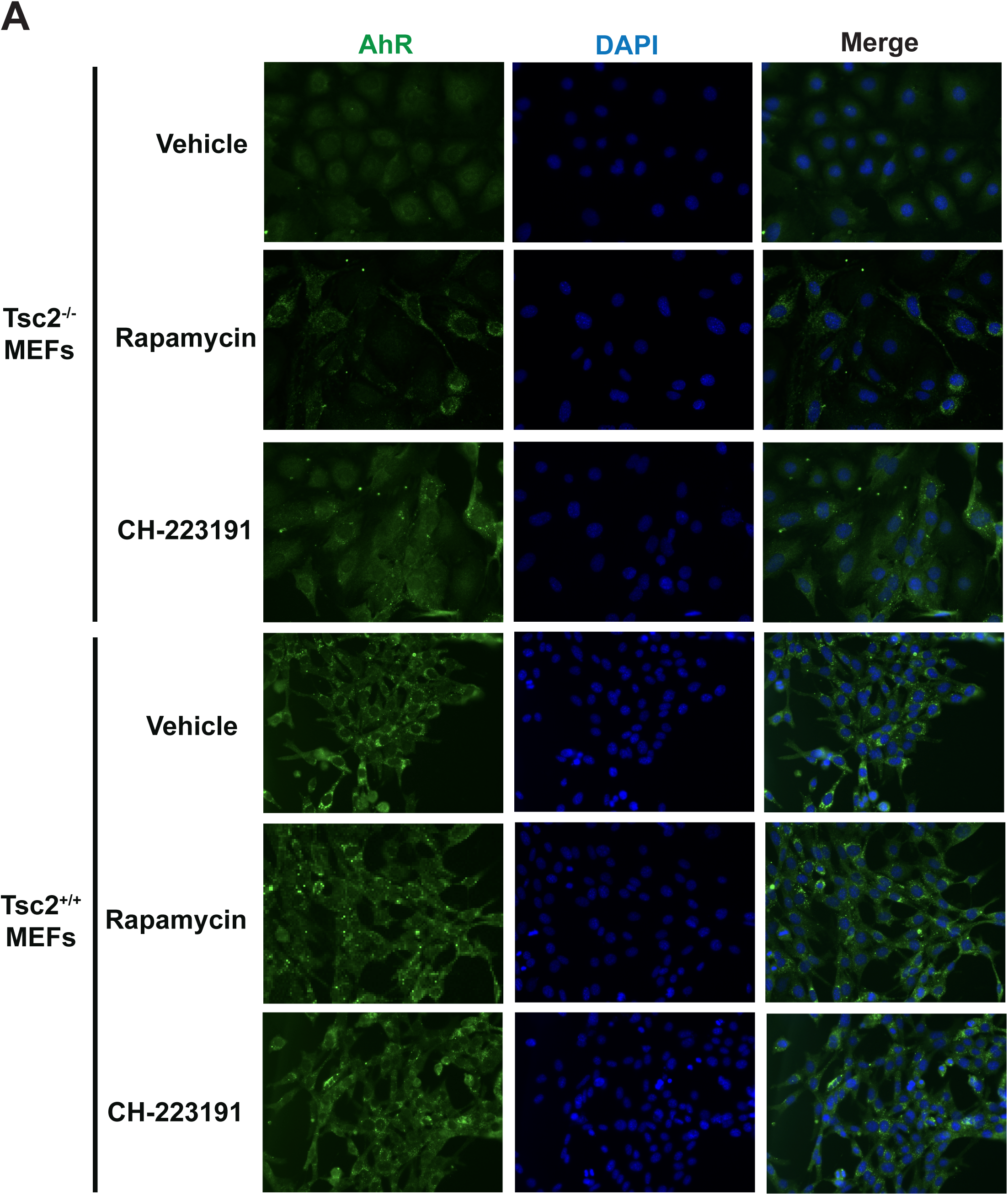
AhR is localized in the nucleus of TSC2-deficient cells. *Tsc2*⁻/⁻ and *Tsc2*⁺/⁺ MEFs were treated with vehicle, rapamycin (20 nM) or CH-223191 (10 µM) for 24 h and stained for AhR (green) and DAPI (blue). AhR co-localized extensively with DAPI in vehicle-treated Tsc2⁻/⁻ MEFs, indicating constitutive nuclear localization, whereas AhR in Tsc2⁺/⁺ MEFs was predominantly cytoplasmic. Rapamycin and CH-223191 each redistributed AhR from the nucleus to the cytoplasm in *Tsc2*⁻/⁻ MEFs, and neither agent altered AhR localization in Tsc2⁺/⁺ MEFs. Representative images from 3 independent experiments, acquired with identical settings; Scale bar, 100µm.

### AhR inhibition induces metabolic reprogramming in TSC2-deficient cells

To define the metabolic consequences of AhR inhibition, we performed steady-state metabolite profiling by LC-MS in *Tsc2*^−/−^ MEFs treated with SR-1 (2 µM, 24 hours) or vehicle. Unsupervised analysis of the top 50 significantly altered metabolites revealed distinct metabolic signatures in SR-1-treated cells compared with vehicle-treated *Tsc2*^−/−^ controls (Fig. 5A). Metabolite set enrichment analysis (MSEA) identified glutamate metabolism as the most significantly enriched pathway among SR-1-regulated metabolites, followed by ammonia recycling, aspartate metabolism, arginine and proline metabolism and the urea cycle. Glutathione metabolism was also enriched (*p* = 0.03; Fig. 5B). The enrichment signature is therefore dominated by amino acid and nitrogen metabolism, consistent with loss of a scavenging route that supplies amino acids. These results link AhR activity to amino acid and antioxidant homeostasis in TSC2-deficient cells.

**Figure 5:**
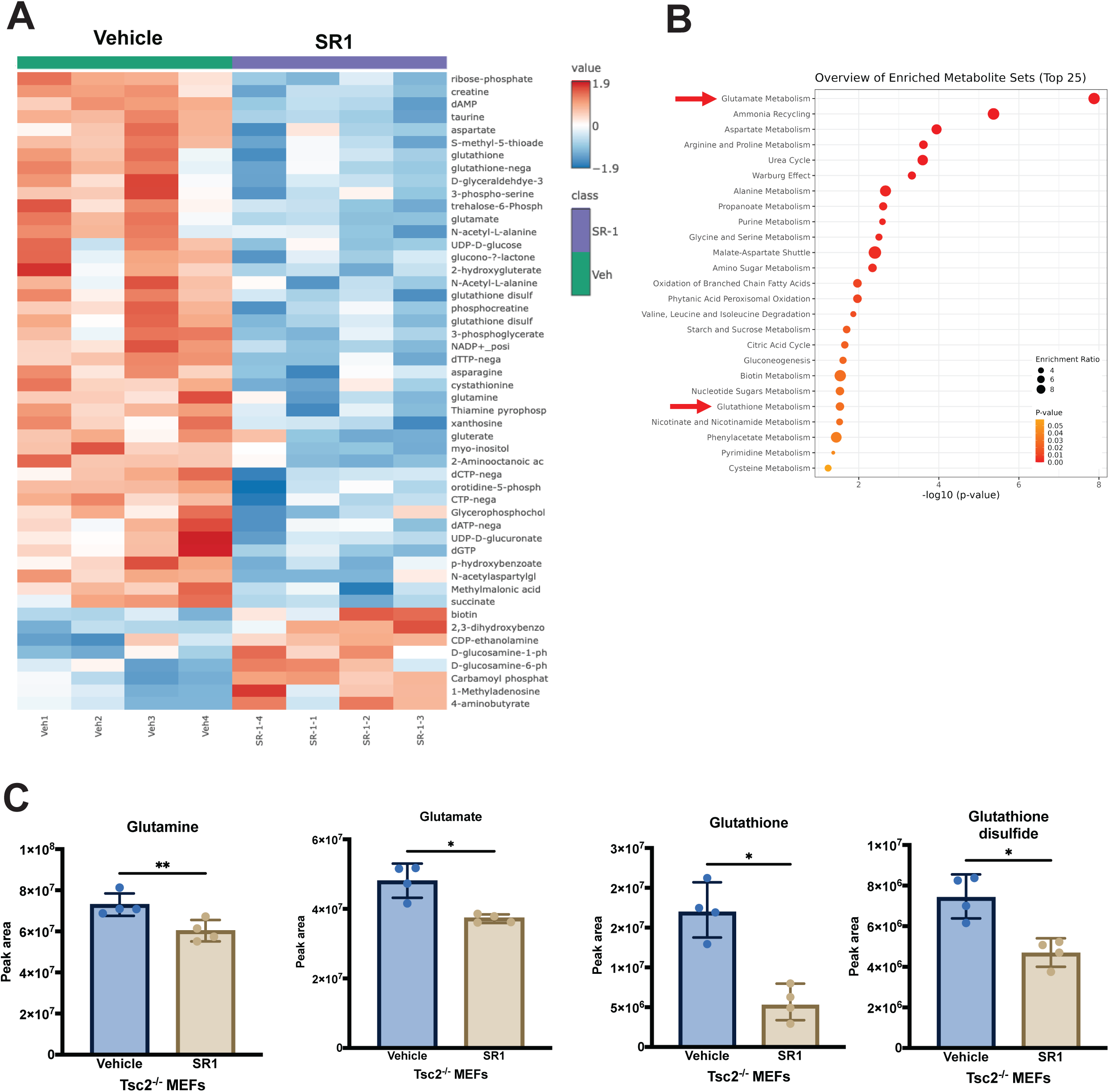
Targeting AhR-mediated macropinocytosis inhibits glutathione metabolism. **(A)** Heatmap of the top 50 significantly altered metabolites (p < 0.05) in *Tsc2*⁻/⁻ MEFs treated with SR-1 (2 µM, 24 h) relative to vehicle. Increased metabolites are shown in red and decreased metabolites in blue; each column is one biological replicate. **(B)** Metabolite set enrichment analysis (MetaboAnalyst 6.0) of SR-1-regulated metabolites. Glutamate metabolism was the most significantly enriched set, followed by ammonia recycling, aspartate metabolism, arginine and proline metabolism and the urea cycle; glutathione metabolism was also enriched (p = 0.03). Arrows mark glutamate and glutathione metabolism. **(C)** Individual metabolites after SR-1 treatment (2 µM, 24 h): glutamine −17% (p < 0.01), glutamate −22% (p < 0.05), reduced glutathione −70% (p < 0.05) and glutathione disulfide −37% (p < 0.05). n = 4 biological replicates per condition; peak areas were normalized to protein concentration. Data are mean ± SD; unpaired two-tailed Student t test. *, p < 0.05; **, p < 0.01.

Analysis of individual metabolites within these pathways revealed that glutamine was decreased by approximately 17% (*p* < 0.01), glutamate by approximately 22% (*p* < 0.05), reduced glutathione (GSH) by approximately 70% (*p* < 0.05), and glutathione disulfide (GSSG) by approximately 37% (*p* < 0.05) following SR-1 treatment (Fig. 5C). Interestingly, the GSH:GSSG ratio was decreased indicating a shift towards a more oxidized glutathione pool. This finding suggests that AhR-dependent macropinocytosis normally delivers extracellular material that supports glutathione biosynthesis, while the reductions in glutamine and glutamate are identifying a deficit in macropinocytic amino acid acquisition.

### Pharmacologic targeting of the kynurenine–AhR axis suppresses tumorigenesis in TSC

We next asked whether the kynurenine pathway can be targeted to suppress TSC tumor growth *in vivo*. We employed a subcutaneous tumor model using TSC2-deficient 105K cells to evaluate the efficacy of AhR antagonists and kynurenine pathway inhibitors in comparison with rapamycin. In an initial efficacy study using the AhR inhibitor SR-1, tumor-bearing mice were treated with SR-1 (6mg/kg, Monday to Friday) by intraperitoneal injection. SR-1 treatment produced an approximately 70% reduction in tumor volume compared with vehicle-treated controls (*p* < 0.0001; n = 8 per group; Fig. 6A). Rapamycin (3 mg/kg, Monday, Wednesday and Friday), the standard of care for TSC-associated tumors, slowed tumor growth by approximately 60% without inducing regression, consistent with its cytostatic activity in mTORC1-driven tumor models (Fig. 6A). Immunohistochemical analysis of tumors revealed that SR-1 treatment decreased Ki-67 staining, a marker of active proliferation, and reduced phosphorylation of S6 kinase (Thr389) levels in treated tumors, indicating reduced proliferation and reduced mTORC1 pathway output *in vivo* (Fig. 6B). Rapamycin-treated tumors similarly showed reduced Ki-67 and phospho-S6 kinase, consistent with the on-target mTORC1 inhibitory activity of this agent (Fig. 6B).

**Figure 6:**
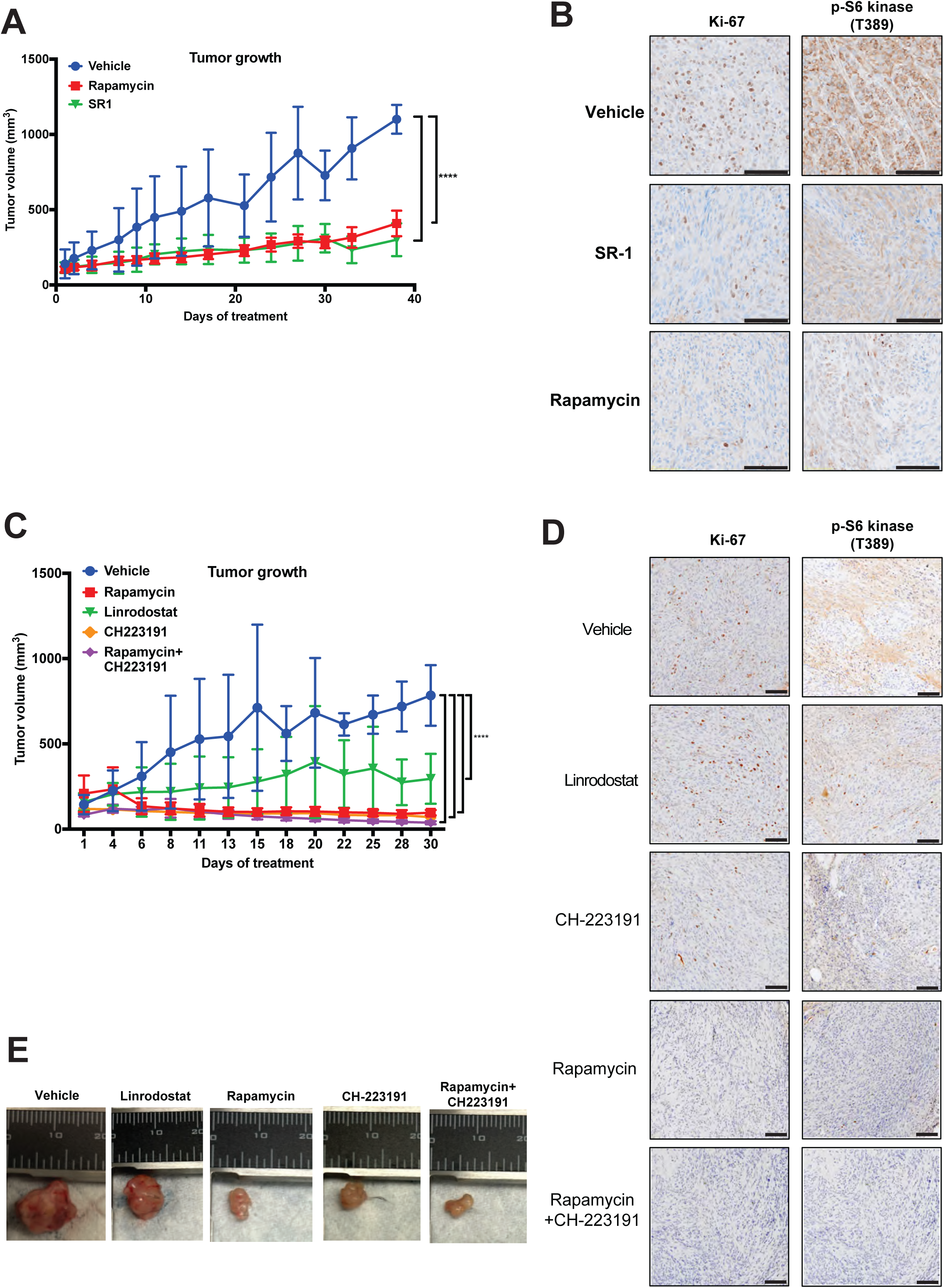
Therapeutic targeting of the kynurenine pathway blocks tumorigenesis in TSC. **(A)** Mice bearing subcutaneous TSC2-deficient 105K tumors were randomized at a mean tumor volume of 100 mm³ to vehicle, SR-1 (6mg/kg intraperitoneally, Monday to Friday) or rapamycin (3 mg/kg, intraperitoneally, Monday, Wednesday and Friday); n = 8 per group. At day 38, SR-1 reduced tumor volume by approximately 70% relative to vehicle (p < 0.0001), and rapamycin slowed tumor growth by approximately 60%. **(B)** Representative Ki-67 and phospho-p70 S6 kinase (Thr389) immunohistochemistry of tumors from the study in A. phospho-p70 S6 kinase and ki67 staining was decreased in the SR1 and rapamycin treated tumors. Scale bar, 100 µm. **(C)** A second cohort was randomized to vehicle, CH-223191 (10 mg/kg, Monday to Friday), linrodostat (10 mg/kg, Monday to Friday), rapamycin (3 mg/kg, Monday, Wednesday and Friday) or rapamycin combined with CH-223191 at the same doses and schedules; n = 8 per group. At day 30, CH-223191 reduced tumor volume by approximately 90% (p < 0.0001), linrodostat by approximately 60% (p < 0.0001) and rapamycin by approximately 90%, and the combination of rapamycin and CH-223191 reduced tumor volume by approximately 95%. **(D)** Representative Ki-67 and phospho-p70 S6 kinase (Thr389) immunohistochemistry of tumors from the study in C. Staining was not quantified. Scale bar, 100 µm. **(E)** Representative images of excised tumors from each treatment arm of the study in C. Data are mean ± 95% confidence interval. Tumor growth curves were analyzed by two-way ANOVA with treatment and time as factors, followed by Dunnett’s multiple comparisons test against the vehicle group; asterisks denote the comparison at the final time point. In C, two pre-specified comparisons were made at the final time point within the same ANOVA, with Bonferroni correction for the two tests: rapamycin plus CH-223191 versus rapamycin alone.

To test a structurally distinct AhR antagonist and to compare AhR blockade with upstream enzyme inhibition, we performed a second study incorporating CH-223191 (10 mg/kg, Monday to Friday), Linrodostat (10 mg/kg, Monday to Friday), and an arm treated with the combination of rapamycin and CH-223191. Surprisingly, CH-223191 treatment resulted in an approximately 90% reduction in tumor volume compared with vehicle-treated controls (*p* < 0.0001; n=8 per group; Fig. 6C). Kynurenine pathway inhibition with Linrodostat suppressed tumor growth by approximately 60% (*p* < 0.0001; Fig. 6C). Rapamycin reduced tumor volume by approximately 90%, and the combination of rapamycin and CH-223191 produced the largest effect observed, an approximately 95% reduction (Fig. 6C). Immunohistochemical analysis of tumors showed reduced Ki-67 and phospho-S6 kinase (Thr389) staining in all four treatment arms relative to vehicle (Fig. 6D). Representative images of excised tumors from each treatment cohort are shown in Fig. 6E. Together, these results provide pharmacologic evidence that the kynurenine–AhR axis is a therapeutically tractable vulnerability in TSC2-deficient tumors and support further evaluation of AhR antagonists, alone or combined with rapalogs, in TSC and LAM.

## Discussion

This study identifies the tryptophan–kynurenine–AhR–macropinocytosis axis as a previously unrecognized metabolic vulnerability in TSC2-deficient cells. Trp alone among the essential amino acids induces macropinocytosis (Fig. 1C) pointing toward a catabolite-mediated rather than a classical amino acid sensing mechanism: the canonical amino acid activators of mTORC1, leucine, arginine and glutamine (30), did not induce macropinocytosis. The decrease of intracellular kynurenine following Trp supplementation (Fig. 1D) is consistent with kynurenine being consumed as rapidly as it is generated, although steady-state pools cannot establish this and isotopic tracing will be required to demonstrate increased flux directly. The relationship between mTORC1 and Trp catabolism appears to be bidirectional: IDO1 has been reported to suppress a Trp sufficiency signal that activates mTOR (27), and our data show the reciprocal arm, in which mTORC1 hyperactivation upregulates *Ido1* and *Tdo2* (Fig. 2B and C) to sustain kynurenine production.

The transcriptional upregulation of *AhR* (approximately 50-fold), *Tdo2* (approximately 45-fold), and *Ido1* (7.5-fold) in *Tsc2*^−/−^ cells, and the strong inhibition of *Ido1* and *Tdo2* expression by rapamycin, identify this as an mTORC1-regulated transcriptional program. The rapamycin sensitivity of *Ido1* and *Tdo2* raises the possibility that rapalogs partially suppress kynurenine pathway activity in TSC patients, and that combining rapalogs with AhR antagonists could achieve more complete blockade, consistent with the greater efficacy of the rapamycin plus CH-223191 combination in our *in vivo* experiments (Fig. 6C). The mechanism linking mTORC1 to *Ido1* and *Tdo2* transcription remains to be determined. However, an autocrine IL-6–STAT3 signal could drive the constitutive IDO1 expression acting in concert with AhR (31). Additionally, TSC2-deficient cells upregulate IL-6 transcription and secretion, with elevated circulating IL-6 in patients with LAM (31).

Our data extend the cell-autonomous pro-tumorigenic functions of the kynurenine–AhR axis described by Opitz and colleagues in glioblastoma (22), adding macropinocytic nutrient scavenging as a downstream effector mechanism in the context of mTORC1 hyperactivation. Constitutive nuclear AhR in *Tsc2*^−/−^ cells (Fig. 4) was reversed by both rapamycin and CH-223191, consistent with an mTORC1-dependent, ligand-driven process. The transcriptional feedback loop whereby AhR sustains *Ido1* and *Tdo2* expression (Fig. 3C) may explain why AhR antagonists were more effective than IDO1 inhibition alone in our *in vivo* studies. Our findings sit alongside the observation that mTORC1 activity suppresses the catabolism of macropinocytosed protein and thereby limits the use of extracellular protein as an amino acid source (14). That constraint does not appear to operate in TSC2-deficient cells: uptake and lysosomal processing of exogenous albumin are both increased in *Tsc2*^−/−^ cells, and inhibition of macropinocytosis reduces lysosomal proteolysis and proliferation (18,32), consistent with the direct demonstration of cell-autonomous catabolism of extracellular protein in tumors *in vivo* (33). Chronic mTORC1 activation in TSC and LAM is compatible with the catabolism of internalized protein, and the Trp-driven increase in uptake reported here is likely to yield free amino acids rather than undegraded cargo. Palm *et al*, and subsequently Nofal *et al*, showed that mTOR activity limits the amino acid yield of internalized protein (34,35). Whether that constraint applies only to growth factor-driven activation, could be resolved by measuring protein catabolic flux directly under Trp supplementation.

The 70% depletion of reduced glutathione following AhR inhibition (Fig. 5C) points to a second, distinct mechanism by which disruption of the kynurenine–AhR axis impairs TSC2-deficient cell survival. Because mTORC1-hyperactive cells generate elevated ROS through increased mitochondrial activity and anabolic biosynthesis (36), they depend on adequate GSH reserves for redox homeostasis, and further compromise of antioxidant capacity is poorly tolerated in this context (37). The observed decrease in glutamine, glutamate, and GSH points to a deficit in macropinocytic amino acid acquisition that cannot be compensated by transporter-mediated uptake under the high biosynthetic demands of TSC2-deficient cells (38). However, the disparity between the small changes in glutamine and glutamate and the large dicrease in GSH suggests that AhR may also act on glutathione metabolism more directly, for example through its documented crosstalk with NRF2 (39). If confirmed, this oxidative vulnerability would represent a mechanism distinct from nutrient deprivation through which AhR antagonism limits the growth of TSC2-deficient tumors.

The approximately 70% and 90% reductions in tumor volume produced by SR-1 and CH-223191, respectively, and the combination of CH-223191 with rapamycin was more effective than either alone (Fig. 6A and C). Recent trials on melanoma using epacadostat (IDO1 inhibitor) (40) shows minimal effect, which is consistent with our observation that IDO1 inhibition alone was less effective than direct AhR antagonism and supports targeting the pathway at the level of AhR rather than at individual upstream enzymes, where redundancy between IDO1 and TDO2 provides a possible route to escape.

Collectively, our data place tryptophan catabolism upstream of macropinocytic nutrient scavenging in TSC2-deficient cells and identify AhR as the node that can be therapeutically targeted.

## Authors’ Contributions

**M. Moyet:** Formal analysis, validation, investigation, visualization methodology, writing–original draft, review and editing. **A. McGrath-Conwell**: Formal analysis, investigation, writing–review and editing. **E. Gray:** Investigation, methodology, writing–review and editing. **S. Jalbert-Jones:** Investigation, methodology, review and editing. **S. Lafleur:** Investigation, writing-review and editing. **C. Rosenberg:** Investigation, review and editing. **V. Patel**: Investigation, methodology, review and editing. **J. Gray:** Investigation, methodology, writing–review and editing. **W. Cox:** Investigation, methodology, review and editing. **E.P. Henske**: Resources, funding acquisition, review and editing. **H. Filippakis**: Conceptualization, resources, formal analysis, supervision, funding acquisition, investigation, visualization, methodology, writing–original draft, writing–review and editing.

## Acknowledgments

This work was supported by NIGMS 5P20GM152330 to H.F. We acknowledge the COBRE Center for Cell Signaling Research for use of the In vitro Analytical Core (IVAC), the COBRE Center for Pain Research for use of the Histology & Imaging Core and Behavior Core at UNE and John Asara, Ph.D. at the BIDMC Mass Spectrometry Core at BIDMC, Boston, MA.

---

**Supplementary Figure S1. Tryptophan, but not kynurenine, supports macropinocytosis in TSC2-deficient cells.**

**(A)** *Tsc2*⁻/⁻ MEFs in amino acid-free medium repleted with Trp (68 μM) or kynurenine (Kyn; 100 μM), with or without linrodostat (20 μM) or EIPA (25 μM), were assayed for 70 kDa FITC-dextran uptake by flow cytometry. Trp increased dextran uptake (p < 0.0001); linrodostat abolished the increase, returning uptake to a level indistinguishable from amino acid-free medium, and EIPA reduced it by approximately 80% (p < 0.0001). Kyn did not increase uptake above the amino acid-free baseline, co-treatment with linrodostat produced no further change, and EIPA reduced uptake in Kyn-supplemented cells by approximately 70% (p< 0.0001), to the same level as amino acid-free cells treated with EIPA. **(B)** Tsc2⁻/⁻ and Tsc2⁺/⁺ MEFs were treated with the LAT1 inhibitor JPH-203 (100 or 200 μM) or EIPA (25 μM). Basal dextran uptake was higher in Tsc2⁻/⁻ than in Tsc2⁺/⁺ cells (p < 0.0001); EIPA reduced uptake in both lines (p < 0.01), whereas JPH-203 had no effect at either concentration. **(C)** Representative images of Tsc2⁻/⁻ and Tsc2⁺/⁺ MEFs in amino acid-free medium with or without Kyn (100 μM) and with or without EIPA (25 μM), showing no increase in dextran-positive macropinosomes following Kyn supplementation in either line. Scale bar, 100 μm. n = 3 biological replicates. Data are mean ± SD; two-way ANOVA with Bonferroni correction. ns, not significant; **, p < 0.01; ****, p < 0.0001.

