## Supplementary figures and images for "Tryptophan catabolism drives AhR-dependent macropinocytosis and tumorigenesis in tuberous sclerosis complex"

### Supplementary Figure 1

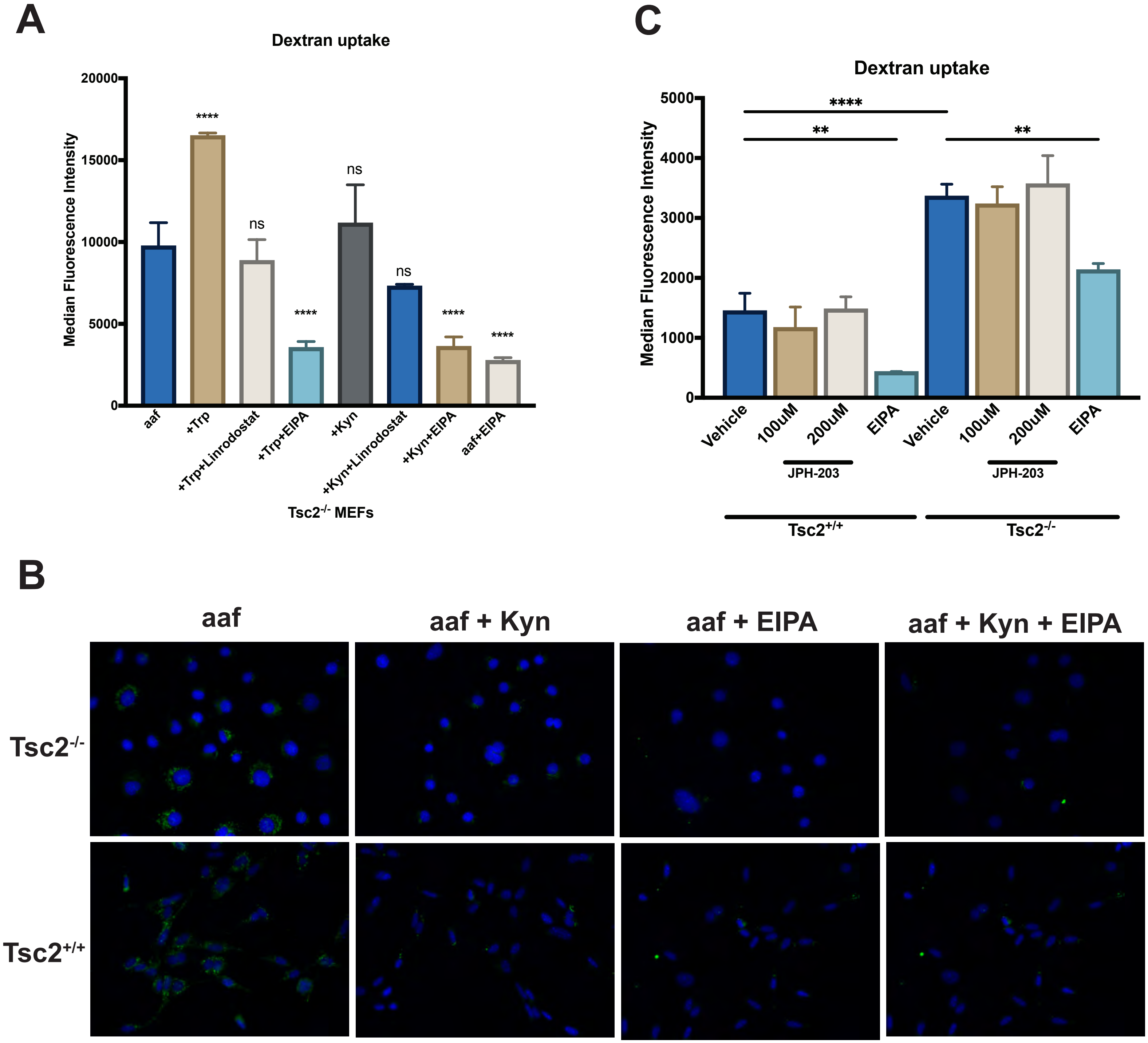


Supplementary figure 1
